# Environmental stress amplifies competitive asymmetry and drives divergent hybrid zone outcomes

**DOI:** 10.1101/2025.10.23.684133

**Authors:** Brynn E. Johnson, Cameron Adkins, Taylor N. Black, Mysia Dye, Isabelle Mendoza, Rachel L. Moran

## Abstract

Community persistence depends on the balance between abiotic constraints and biotic interactions. Environmental stress can either sort species by physiological limits or amplify competitive asymmetries, producing coexistence, exclusion, or collapse. We tested these alternatives in two replicate hybrid swarms between orangethroat and orangebelly darters (*Etheostoma pulchellum* and *E. radiosum* spp. complex) with contrasting outcomes: long-term coexistence in the Blue River versus collapse in the Washita River. We combined critical-thermal-maximum (CT_max_) assays with standardized feeding experiments to evaluate physiological tolerance, competitive exclusion, and stress-amplified competition. CT_max_ varied with river, sex, and body size but not consistently between species, indicating that local history and demography outweighed intrinsic physiological differences. In contrast, competition trials revealed strong, temperature-dependent asymmetries: *E. pulchellum* dominated in the cooler, stable Blue River, whereas *E. radiosum* spp. gained a foraging advantage under high temperatures in the warmer Washita River drainage. These results support the prediction that abiotic stress amplifies competitive asymmetries, flipping dominance and explaining divergent hybrid zone outcomes. More broadly, our study links hybrid zone dynamics to coexistence theory, showing that climate extremes can shift competitive balance and determine whether secondary contact results in persistence or loss.

## Introduction

The persistence of ecological communities depends on the balance between abiotic filtering and biotic interactions. When diverged species come into secondary contact, this balance determines whether they coexist, hybridize, or one excludes the other. Abiotic filtering occurs when environmental conditions such as temperature or oxygen availability exclude individuals unable to tolerate them (Dunson & Travis 1991, Kraft et al. 2015). In many cases, such filtering operates through differences in physiological tolerance, including differential survival or performance under abiotic stress (Lee et al. 2003). Biotic interactions, in contrast, determine persistence through mechanisms such as competition or predation. Classic coexistence theory predicts that environmental stress can either sort species by tolerance or amplify competitive asymmetries, tipping communities toward exclusion or collapse (Barton & Hewitt 1985, Seehausen 2006, Chesson 2000). Distinguishing between these pathways is essential for predicting how community composition and species ranges shift under changing environments.

Abiotic stressors such as heat and hypoxia can structure communities by filtering species according to physiological limits (Sunday et al. 2012, Payne et al. 2016). By contrast, biotic interactions such as resource competition can drive exclusion even in the absence of strong physiological asymmetries (Chesson 2000). In this study we focus on exploitative competition, in which individuals indirectly compete by depleting shared resources rather than through direct aggression (Tilman 2007). Most studies focus on one axis or the other, but anthropogenic disturbance simultaneously alters abiotic and biotic environments, and their combined effects on persistence remain poorly understood.

Freshwater fishes are particularly vulnerable to environmental change due to their limited dispersal abilities and sensitivity to fluctuations in temperature and hydrology (Heino et al. 2009, Bush & Hoskins 2017). Habitat alteration increasingly forces previously isolated species into secondary contact, where hybridization can generate long-lasting hybrid swarms (Seehausen et al. 2008, Porretta & Canestrelli 2023). These hybrid systems provide natural experiments for examining how physiological tolerance and competitive dynamics interact to determine persistence after contact.

Darters (Percidae) provide an excellent system to investigate the mechanisms that promote and impede species diversification (Mendelson 2003, Mendelson et al. 2007, Keck & Near 2009, Zhou & Fuller 2016, Moran et al. 2017, Moran & Fuller 2018). Darters are one of the most diverse vertebrate radiations in North America, with over 250 described species distributed across the eastern half of the United States (Page & Burr 2011). Yet, darters are also disproportionately affected by anthropogenic habitat disturbances, with over 50% of species in the clade considered imperiled (Helfman et al. 2009). Two well-documented, replicate hybrid swarms between the plains orangethroat darter (*Etheostoma pulchellum*) and members of the orangebelly darter complex (*Etheostoma radiosum* spp.) in the Blue River and Washita River drainages in southeastern Oklahoma provide an exciting opportunity to study the behavioral and environmental factors influencing species dynamics in a relatively isolated system (Branson & Campbell 1969, Echelle et al. 1974, Matthews et al. 2016). In the Blue River, *E. pulchellum, E. radiosum cyanorum*, and their hybrids have been present for >80 years. Conversely, both *E. pulchellum* and *E. radiosum paludosum* were documented as co-occurring and hybridizing in Little Glasses Creek (Washita River drainage) in 1985, but since 2003 only *E. radiosum paludosum* has been observed in this previous hybrid zone (Matthews et al. 2016). This indicates local extirpation of *E. pulchellum* from Little Glasses Creek. These contrasting outcomes for the hybrid swarms within the Blue River and Washita River drainages allow a direct test of whether physiological tolerance or exploitative competition better predicts persistence after secondary contact.

Here we quantify (i) thermal limits (critical thermal maximum, CT_max_) for each species in both sympatric hybrid zone populations and allopatric populations, and (ii) exploitative competition via standardized feeding assays within and between species from sympatric populations to assess the respective contributions of physiological capability and competitive interactions to the long-term stability of the Blue River hybrid zone and collapse of the Little Glasses Creek hybrid zone. It has been hypothesized that the disappearance of *E. pulchellum* from Little Glasses Creek could result from environmental filtering (Matthews et al. 2016), where only those individuals or species with sufficient tolerance to high temperatures and low oxygen survive. Such filtering could act through differences in physiological tolerance (i.e., differential survival under thermal or oxygen stress). In support of this, darters show seasonal and population-level variation in CT_max_ in response to ambient temperatures (Hlohowskyj & Wissing 1985, Firth et al. 2021), and comparative fish work indicates that species’ upper thermal tolerances correlate with environmental temperature regimes (Comte & Olden 2017). Complementing this filtering view, trait-based competition models demonstrate how environmental constraints and interspecific interactions jointly shape community composition (Wong et al. 2025).

From this framework, we derive three explicit predictions about the mechanisms driving species persistence after secondary contact. If physiological tolerance explains the observed pattern, we expect members of the *E. radiosum* spp. complex to exhibit higher CT_max_ than *E. pulchellum* in both sympatry and allopatry. Alternatively, if competitive exclusion is the primary driver, we predict little difference in CT_max_ between species but higher foraging efficiency and stronger resource depletion by members of the *E. radiosum* spp. complex in mixed-species feeding assays. Finally, if abiotic stress amplifies competitive asymmetries, we expect interspecific competition to strengthen under high temperature or low oxygen. By testing these predictions in replicate hybrid swarms, our study links classic hybrid zone theory with eco-physiological mechanisms, advancing a general framework for predicting whether secondary contact produces coexistence, exclusion, or collapse (Sunday et al. 2012, Dressler 2023).

## Methods

*Etheostoma pulchellum* and *E. radiosum* spp. were collected by kick-seine in February 2024 and 2025 from Oklahoma, Texas, and Arkansas and transported back to the lab in aerated coolers within two days of collection. Sympatric *E. pulchellum* were collected from the Blue River hybrid swarm site, under the Connorville Hwy 99 bridge (34°27’15.8”N 96°38’08.9”W) and Pennington Creek (Washita River drainage) in close proximity to the Tishomingo National Fish Hatchery (34°21’04.9”N 96°42’37.1”W) (Figure 1). Sympatric *E. radiosum cyanorum* were collected from the Blue River hybrid swarm site and *E. radiosum paludosum* were collected from the Washita River hybrid swarm site, located in Little Glasses Creek (34°01’32.4”N 96°41’53.8”W) (Figure 1). Allopatric *E. pulchellum* were collected from Salado Creek in the Brazos River Drainage at 30°56’44.6”N 97°32’00.5”W, located in Pace Park (Salado, TX). Finally, allopatric *E. radiosum radiosum* were collected from Collier Creek (Caddo River drainage) in Caddo Gap, AR at 34°24’51.9”N 93°37’50.6”W. Fish were allowed to acclimate in 37.9 L tanks separated by species and sex with a 12:12 light:dark cycle at ambient room temperature (15.6°C) and were fed frozen bloodworms daily *ad libitum*.

**Figure 1.**
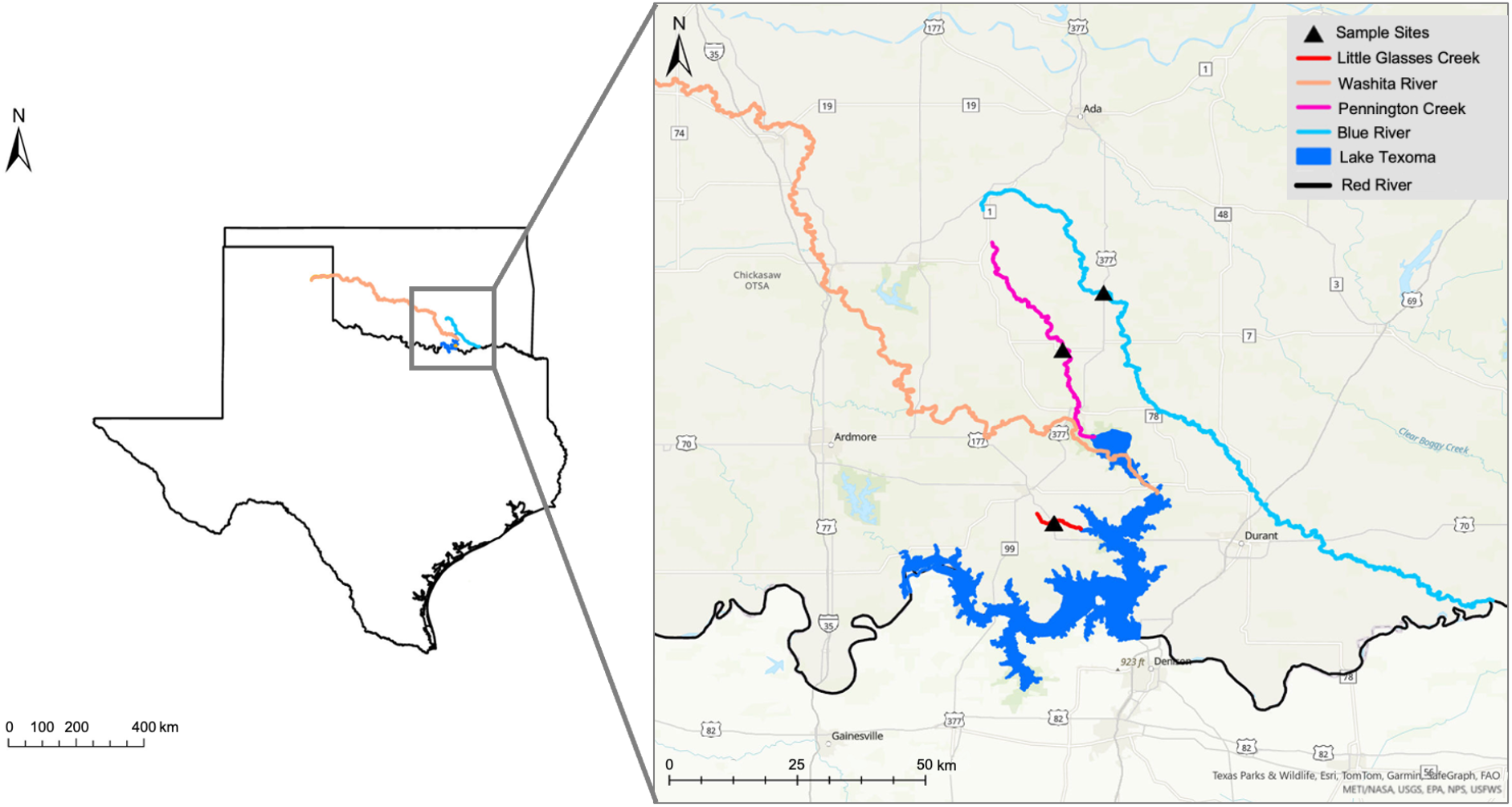
Map of *E. pulchellum* x *E. radiosum* spp. hybrid zone sampling locations in southeastern Oklahoma showing the Washita River (orange) and Blue River (light blue) in relation to Lake Texoma (dark blue). Sampling sites are indicated by black triangles and include the main stem of the Blue River and two tributaries of the Washita River: Pennington Creek (pink) and Little Glasses Creek (red). The Red River forms the Oklahoma-Texas border, with Oklahoma to the north and Texas to the south. Inset map shows the location of these drainages within Texas and Oklahoma. Scale bars in kilometers (km). The map was created in ArcGIS Pro 3.0.0 using Esri World Topographic Map basemap. Stream and river shapefiles were obtained from the Oklahoma Water Resources Board (for Oklahoma streams and major rivers) and the U.S. Geological Survey (for Texas rivers).

### Critical Thermal Maximum (CT_max_) Assays

CT_max_ trials were conducted in June 2024 and 2025 after fish were no longer in spawning condition (females non-gravid, males dull in coloration). Prior to trials, individuals were held for 1–3 months at ambient room temperature (16 ± 1 °C) in aerated 37.9 L tanks. Following Oliveira et al. (2021), fish were placed in an insulated 68 L Igloo cooler containing 8 L of water at 16 ± 1 °C. Eight perforated coffee-filter containers, affixed to the cooler bottom with aquarium-safe sealant, held individual fish during trials. Dissolved oxygen was maintained with an aeration stone, and water was mixed with a Rio Plus 90 Aqua Pump (2.8 W, TAAM Inc.) to ensure temperature homogeneity. Heating coils connected to a voltage transformer raised the temperature at a controlled rate of 0.3 °C min^-1^, monitored with a YSI Pro 20 probe.

Loss of equilibrium (LOE) was defined as rapid muscle spasms followed by loss of the ability to remain upright (Lutterschmidt & Hutchison 1997). At LOE, fish were immediately transferred to a 19 L (5 gal) recovery tank at 25 °C containing aeration and API Stresscoat (5 mL). Water was replaced between trials. A total of 129 individuals were assayed (Table 1), with length and weight recorded prior to testing.

**Table 1.** Focal species, drainage, geography (sympatric or allopatric), and sample size (by sex) for CT_max_ trials.

| Species name | Drainage | Geography | *n* (male, female) |
| --- | --- | --- | --- |
| *E. pulchellum* | Washita | Sympatric | 7,13 |
| *E. radiosum paludosum* | Washita | Sympatric | 17,12 |
| *E. pulchellum* | Blue | Sympatric | 10,10 |
| *E. radiosum cyanorum* | Blue | Sympatric | 10,10 |
| *E. pulchellum* | Brazos | Allopatric | 9,11 |
| *E. radiosum radiosum* | Caddo | Allopatric | 10,10 |

CT_max_ was analyzed using linear models in R v4.4.1. We fit models with CT_max_ as the response and included species group, river, sex, and body length (centered) as predictors. For sympatric populations (Blue and Washita rivers), we tested for species differences and species x river interactions using Type III ANOVA in the *car* package. We also ran a broader model including all four rivers (Blue, Washita, Brazos, Caddo) with species and river as additive fixed effects, to assess general geographic variation in CT_max_. Estimated marginal means from the *emmeans* package were used to obtain pairwise contrasts between species within each river, with Tukey adjustments for multiple comparisons.

During the CT_max_ trials, one *E. pulchellum* male from the Washita River displayed an unusually low value that was far outside the range of all other observations. Because this single outlier disproportionately affected species contrasts within the Washita River, we excluded it from analyses focused on sympatric populations to avoid misleading inference. For the broader four-river analysis, however, we retained the individual since it did not influence overall geographic patterns or change the main conclusions. Results with and without the outlier are presented in the Supplementary Materials to ensure transparency.

### Exploitative Competition Trials

We conducted feeding trials to test whether exploitative competition contributed to the collapse of the Washita River hybrid swarm and the persistence of orangebelly darters (*E. radiosum* spp.) in Little Glasses Creek. Orangethroat (*E. pulchellum*) and orangebelly darters from the Blue and Washita River drainages were acclimated for 10 days in aerated 37.9 L tanks, grouped by population and sex, at one of three temperatures (15.5°C, 25.5°C, or 30°C, hereafter “Cold”, “Room”, and “Hot”). Individuals were marked with elastomer tags (Northwest Marine Technology, Inc.) to allow identification, and total length and weight were measured prior to trials.

Following acclimation, fish were maintained without feeding for 72 h prior to standardize digestive state prior to assays (Carmona-Catot et al. 2013, Raby et al. 2025). This fasting interval lies within the range commonly adopted for small benthic fishes and, based on observations in similar experimental systems, does not typically induce measurable stress or significant mass loss (e.g. Killen et al. 2014a; B. Johnson, personal observation). For each trial, one *E. pulchellum* and one *E. radiosum* spp. darter were size-matched (within 5 mm total length) from the same acclimation temperature. Each pair was introduced to a 9.5 L test tank filled with water at their acclimation temperature. After a 5 min acclimation period in the test tank, five bloodworms were simultaneously introduced. We recorded the identity of the “First Worm Winner” (the individual consuming the first worm) and the “Overall Worm Winner” (the individual consuming ≥3 of 5 worms). Trials were replicated with random pairings across all three temperatures.

We analyzed “First Worm Winner” with a logistic mixed model (glmmTMB) using *E. radiosum* spp. victory (yes/no) as the response, fixed effects of temperature (Cold/Room/Hot), river (Blue/Washita), their interaction, pair sex, and species-specific size advantage (*E. radiosum* spp. length − *E. pulchellum* length; centered), and a random intercept for trial.

For “Overall Worm Winner”, we modeled counts of worms eaten (0-5) per individual using a binomial mixed model with fixed effects of species, temperature, river, all interactions, pair sex, and centered length, and a random intercept for trial; trials with both counts missing (n=13) were excluded. Type III Wald tests used sum-to-zero contrasts, and we report odds ratios with 95% CIs and model-adjusted means (EMMs) for inference and visualization.

## Results

### CT_max_ Assays

Our analysis of CT_max_ across sympatric populations detected a significant species x river interaction (Table 2; *F*_1,81_ = 16.3, *p* < 0.001), indicating that relative thermal tolerance differed between rivers. In the Blue River, *E. r. cyanorum* tended to tolerate higher temperatures than *E. pulchellum* (mean difference = 0.60 °C, *t* = –2.44, *p* = 0.017), whereas in the Washita River the pattern was reversed, with *E. pulchellum* exhibiting higher CTmax than *E. r. paludosum* (mean difference = 0.78 °C, *t* = 3.37, *p* = 0.001; Figure 2).

**Table 2.** Type III ANOVA for critical thermal maximum (CT_max_) in sympatric populations of *E. pulchellum* and *E. radiosum* spp.

| Effect | df | F | p-value |
| --- | --- | --- | --- |
| Species | 1 | 0.31 | 0.58 |
| River | 1 | 12.21 | **0.0008** |
| Sex | 1 | 0.2 | 0.657 |
| Length (centered) | 1 | 0.49 | 0.487 |
| Species x River | 1 | 16.3 | **0.0001** |

Note: Washita male *E. pulchellum* outlier excluded. Bolding indicates significance at $p < 0.05$ .

**Figure 2.**
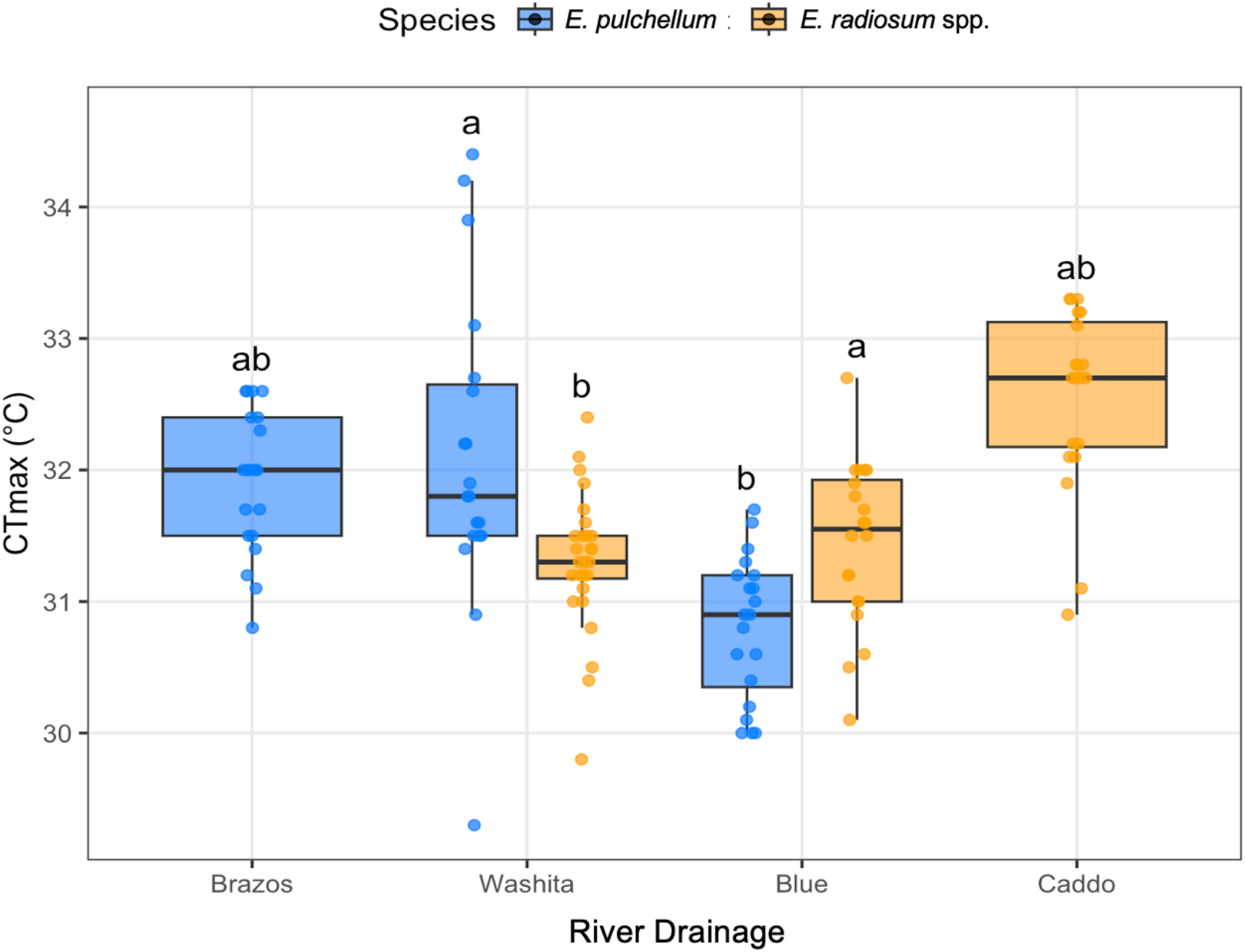
Critical thermal maximum (CT_max_) of *E. pulchellum* (blue) and *E. radiosum* spp. (orange) from sympatric (Blue River, OK; Washita River, OK) and allopatric (Brazos River, TX; Caddo River, AR) populations. Boxplots show medians, interquartile ranges, and ranges, with individual data points overlaid. Letters denote results of post hoc comparisons across all population x species groups; groups sharing a letter are not significantly different (p > 0.05).

Models including allopatric populations revealed strong river effects on CTmax (F_3,121_ = 13.6, p < 0.0001) but no overall species differences (Table S1). In this broader model, CTmax also varied by sex (F_1,121_ = 8.23, p = 0.0049), with females showing slightly higher tolerance (Figure S1), and by body length (F_1,121_ = 5.32, p = 0.023), with larger individuals generally tolerating higher temperatures. Across rivers, CTmax was highest in Caddo River *E. r. radiosum* darters and lowest in Blue River *E. pulchellum* darters (Figure 2).

### Exploitative Competition Trials

We observed a significant effect of river on first-worm capture (mixed-effects logistic regression; OR = 0.54, 95% CI: 0.32–0.93, p = 0.025, Table S2), with *E. radiosum* spp. less likely to capture the first worm in the Washita River than in the Blue River. Specifically, *E. radiosum* spp. won the first worm in ∼40% of Washita trials vs. ∼65% of Blue River trials. Temperature, sex, and relative size did not significantly affect first-worm capture success (all p > 0.1) (Table S2).

Mixed-effects models revealed strong species differences in overall worm consumption that varied with river and temperature (Table 3, Figure 3), with *E. pulchellum* generally consuming more prey in the Blue River, but this advantage reversing at high temperature in the Washita. Across all trials, species identity was a significant predictor of foraging success (χ^2^ = 55.9, p < 0.001), with body length also contributing positively (χ^2^ = 9.2, p = 0.002). Importantly, interactions between species and river (χ^2^ = 74.6, p < 0.001), species and temperature (χ^2^ = 8.2, p = 0.016), and their three-way interaction (χ^2^ = 25.8, p < 0.001) indicate that competitive outcomes depended on both environmental conditions and drainage.

**Table 3.** Type III Wald χ^2^ tests for overall worm consumption (“Overall Worm Winner”) in exploitative competition trials.

| Predictor | $\chi^2$ | df | p-value |
| --- | --- | --- | --- |
| Species | 55.94 | 1 | **<0.001** |
| Temp | 0.08 | 2 | 0.962 |
| River | 0.03 | 1 | 0.859 |
| Sex | 0.21 | 1 | 0.646 |
| Length | 9.18 | 1 | **0.002** |
| Species x Temp | 8.24 | 2 | **0.016** |
| Species x River | 74.6 | 1 | **<0.001** |
| Temp x River | 0.15 | 2 | 0.929 |
| Species x Temp x River | 25.77 | 2 | **<0.001** |

Note: Bolding indicates significance at $p < 0.05$ .

**Figure 3.**
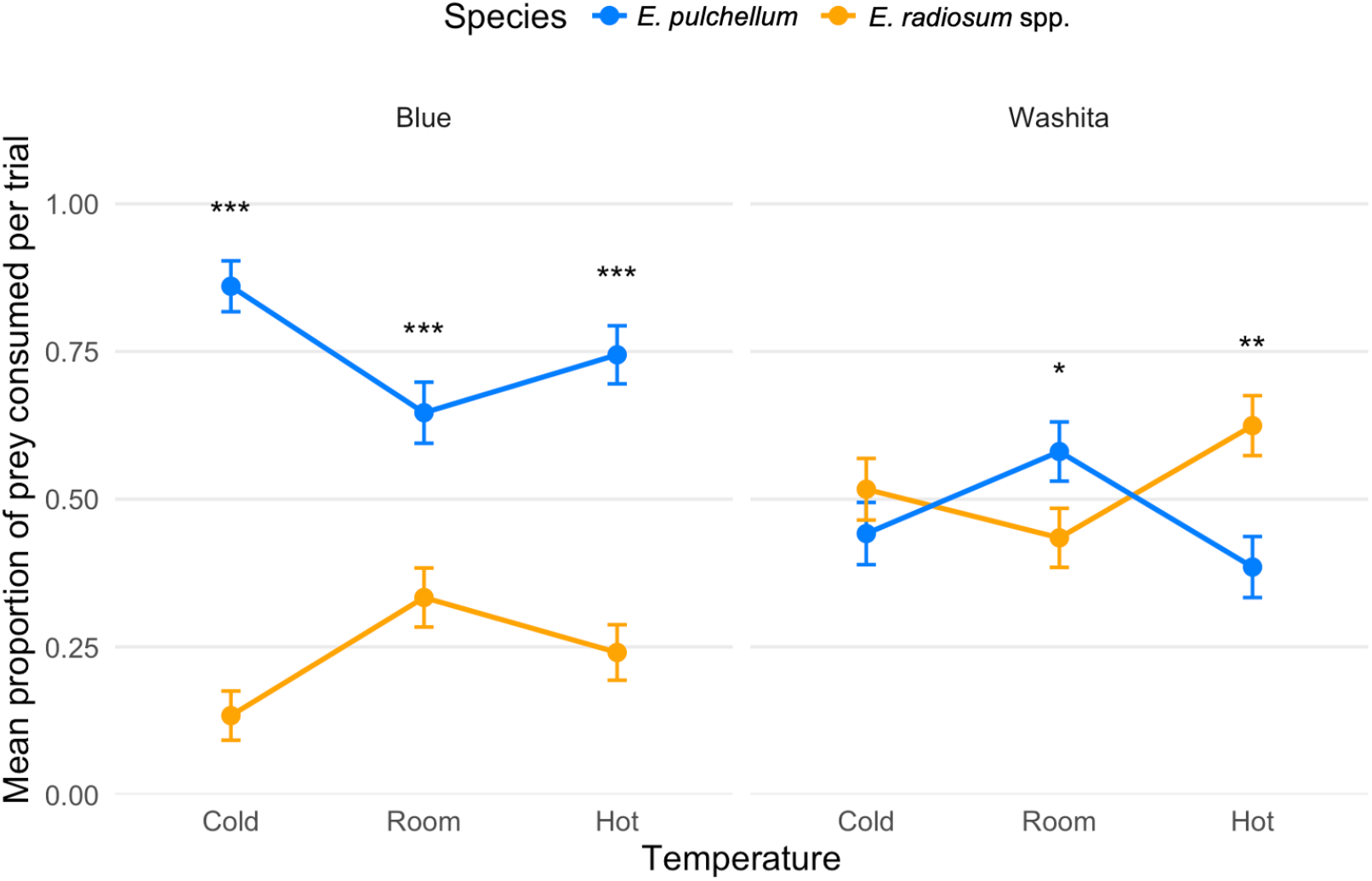
Exploitative competition outcomes between *E. radiosum* spp. (orange) and *E. pulchellum* (blue) darters across two sympatric rivers and three temperature treatments. Points indicate model-adjusted mean proportion of prey consumed per trial (± SE) from binomial mixed-effects models, with lines connecting means across temperatures within species for visualization. Asterisks above comparisons denote significant differences between species within each river x temperature combination (\**p* < 0.05, ** *p* < 0.01, \*\*\**p* < 0.001).

Estimated marginal means clarified these patterns. In the Blue River, *E. pulchellum* consistently dominated, consuming the majority of prey items across temperatures (e.g., OR = 40.0 at Cold, 9.2 at Hot; all p < 0.001; Table S3). By contrast, outcomes in the Washita River shifted toward parity or *E. radiosum* spp. advantage. At Cold and Room temperatures, neither species consistently outcompeted the other (p ≥ 0.05), but at Hot temperatures *E. radiosum* spp. significantly outperformed *E. pulchellum* (OR = 0.38, p = 0.0016) (Figure 3).

## Discussion

Species persistence after secondary contact depends on how abiotic constraints interact with biotic processes such as competition. Our results demonstrate that the contrasting outcomes of two replicate hybrid zones between *E. pulchellum* and members of the *E. radiosum* spp. complex cannot be explained by differences in physiological tolerance alone. Our CT_max_ assays revealed no consistent species-level differences in acute thermal tolerance; instead, species differences were contingent on drainage, and geographic context explained more variation than species identity. These findings suggest that thermal limits are not fixed traits of the lineages but are shaped by local environmental history or demographic context. In the broader four-river comparison, sex and body size also influenced CT_max_, with females and larger individuals generally tolerating higher temperatures. While these effects were modest, they highlight the potential for within-population variation to shape responses to environmental stress, consistent with theory predicting that demographic composition can modulate species’ realized thermal niches (Chesson 2000; Angilletta et al. 2010).

In contrast, the foraging competition assays revealed clear competitive asymmetries that shifted across rivers and thermal environments. *Etheostoma pulchellum* consistently dominated feeding in the Blue River, regardless of temperature, supporting a baseline competitive advantage for this species in that system. In the Washita River, however, competitive outcomes were strongly temperature-dependent: at cooler and intermediate temperatures, neither species dominated, but under elevated temperatures, *E. radiosum paludosum* gained a marked foraging advantage. This pattern aligns with our predictions for competitive exclusion and abiotic stress amplification. While CT_max_ results indicate that both species can physiologically survive acute high temperatures, competition trials show that warming alters the balance of competitive dominance, tipping the outcome toward *E. radiosum* spp. in the Washita system.

The contrasting competition patterns between the Blue and Washita rivers likely reflect long-term environmental and demographic differences between drainages. The Blue River is spring-fed, cooler, and hydrologically more stable, whereas the Washita experiences stronger seasonal fluctuations, periodic drying, and higher summer maxima (Tortorelli 2008; Shivers & Andrews 2013). These baseline differences may modulate energy budgets, foraging pressure, and species interactions. In a stable, cool system like the Blue River, behavioral dominance or foraging efficiency may tip in *E. pulchellum*’s favor. In contrast, episodic thermal and oxygen stress in the Washita River may impose metabolic constraints that reduce the performance of more aggressive species, shifting competitive advantage toward *E. radiosum* spp. Although both species tolerate similar acute heat loads, temperature-dependent variation in metabolism, activity, and risk tolerance can alter competitive hierarchies under stress. In coral reef fishes, such context-dependent dominance has been linked to variation in metabolic rate and temperature-sensitivity of behavior (Killen et al. 2014b, Warren & McCormick 2019, Bailey et al. 2022). More broadly, controlled community experiments in microbes and insects demonstrate that warming and temporal variability can reorganize competitive dominance even without fixed physiological differences (Jiang & Morin 2004, O’Neal & Juliano 2012). Comparable context-dependent shifts occur in amphibians, where thermoregulation and competition interactively determine performance and persistence under warming and habitat modification (Gvoždík 2022; Nowakowski et al. 2018). Together, these parallels suggest that environmental stress can reconfigure competitive hierarchies across taxa whenever species differ subtly in their performance curves, consistent with coexistence theory predicting that stress can flip the sign of competitive asymmetry (Chesson 2000).

The *E. radiosum* complex itself is taxonomically structured, with recent work recommending elevation of *E. radiosum cyanorum* to full species status (*E. cyanorum*) and treatment of *E. radiosum paludosum* as a subspecies of uncertain rank (Matthews & Turner 2019). Divergence within this complex may contribute to the river-specific patterns observed here. Likewise, the taxonomic and genetic diversity of *E. pulchellum* remains unresolved. Preliminary genomic data indicate that the Blue River population is deeply differentiated from those in the Washita and Brazos drainages (Kim et al., in prep.), suggesting long-term isolation and potential local adaptation within *E. pulchellum* itself.

More broadly, the sex and size effects observed in our CT_max_ assays suggest that demographic composition may further modulate persistence under climate stress. Populations skewed toward males or composed of smaller individuals could be especially vulnerable to warming, both because of lower thermal tolerance (Vallin et al. 2025) and because warming may exacerbate the metabolic constraint in larger individuals (Kraskura et al. 2023). These demographic sensitivities imply that conservation strategies should not rely solely on species-level averages; instead, they must account for how local context, environmental extremes, and variation in population structure interact to shape persistence.

Headwater prairie streams frequently become shallow, isolated pools during drought, with high temperature and low oxygen (Matthews 1988). In these conditions, persistence hinges not just on tolerance but on winning scarce resources, so warming can tip competitive balance and determine which lineage persists after secondary contact. This system-specific mechanism explains the Blue-Washita contrast and sets expectations for other intermittency-prone drainages.

This framework has broad implications for understanding range boundaries and predicting biodiversity responses to global change. Thermal limits have long been recognized as key determinants of species distributions (Sunday et al. 2012; Polato et al. 2018), but growing evidence indicates that indirect effects of climate, e.g., through altered species interactions, are equally important (Araújo & Luoto 2007; Cahill et al. 2013). By linking CT_max_ assays with competition experiments, we show that climate extremes can alter competitive hierarchies, triggering local extirpation even when both species can physiologically tolerate high temperatures. This mechanistic insight helps explain why hybrid zones are particularly dynamic under climate change: they represent systems where small shifts in abiotic stress can cascade into large changes in competitive dominance and hybrid persistence.

Our study underscores the value of hybrid swarms as natural experiments for testing coexistence theory. Because hybrid zones integrate gene flow, selection, and ecological sorting, our findings suggest that even subtle thermal differences can cascade into shifts in hybrid swarm stability. Similar eco-physiological feedback could underlie speciation reversal observed in other taxa under climate stress (Seehausen et al. 2008). By explicitly contrasting physiological tolerance with competition under variable abiotic stress, we show that community persistence after secondary contact depends not only on species’ tolerance limits but on how environmental stress reshapes the balance of competitive dominance. This work integrates classical models of hybrid zone dynamics with eco-physiological mechanisms, advancing a general framework for predicting when secondary contact leads to coexistence, exclusion, or collapse. More broadly, our findings highlight how climate extremes can interact with species interactions to drive abrupt shifts in community composition, providing a tractable model for linking evolutionary theory with contemporary biodiversity change.

## Acknowledgements

We thank Kira Delmore, Joshuah Perkin, Daemin Kim, Wynne Radcliffe, Adheeta Dongre, and Gabrielle Welsh for helpful comments on this manuscript. The treatment of animals used in this study followed Texas A&M University’s Institutional Animal Care and Use Committee (IACUC) under AUP #2023-0007. Funding was provided by NSF IOS #2338043 to R.L.M. and by a pilot grant to B.E.J. from the *Integrating Organismal Biology into NEON* Research Coordination Network (NSF RCN #2110233). Fish were collected under Texas Parks and Wildlife Department permit SPR-0323-023, Oklahoma Department of Wildlife Conservation license 10775512, and Arkansas Game and Fish Commission permit 20620232 to R.L.M.

## Data and Code Availability

All data and R scripts will be available from the Dryad Digital Repository (publicly available upon acceptance).

