## Supplementary Materials for "Environmental stress amplifies competitive asymmetry and drives divergent hybrid zone outcomes"

**Supplemental Materials**

In the sympatric rivers analysis (Blue + Washita), there was no overall species effect on CTmax (F_1,82_ = 0.06, p = 0.81), but a significant Species x River interaction (F_1,82_ = 5.74, p = 0.019). This means species differences in thermal tolerance depended on river. Estimated marginal means showed that in the Blue River, *E. pulchellum* and *E. radiosum* *cyanorum* had overlapping CTmax values (31.0 vs. 31.5 °C, p = 0.073). In contrast, in the Washita River, *E. pulchellum* tended to have slightly higher CTmax (31.9 °C) than orangebellies (31.4 °C), though this difference was not significant at the 0.05 level (p = 0.11). In the sensitivity analysis (excluding the extreme low CTmax outlier in a Washita male), the Species x River interaction became stronger (F_1,81_ = 16.3, p < 0.001). Here, *E. pulchellum* had significantly higher CTmax in the Washita (+0.78 °C, p = 0.0011), whereas *E. radiosum* *cyanorum* had higher CTmax in the Blue (+0.60 °C, p = 0.017). Thus, the thermal advantage flipped across drainages.

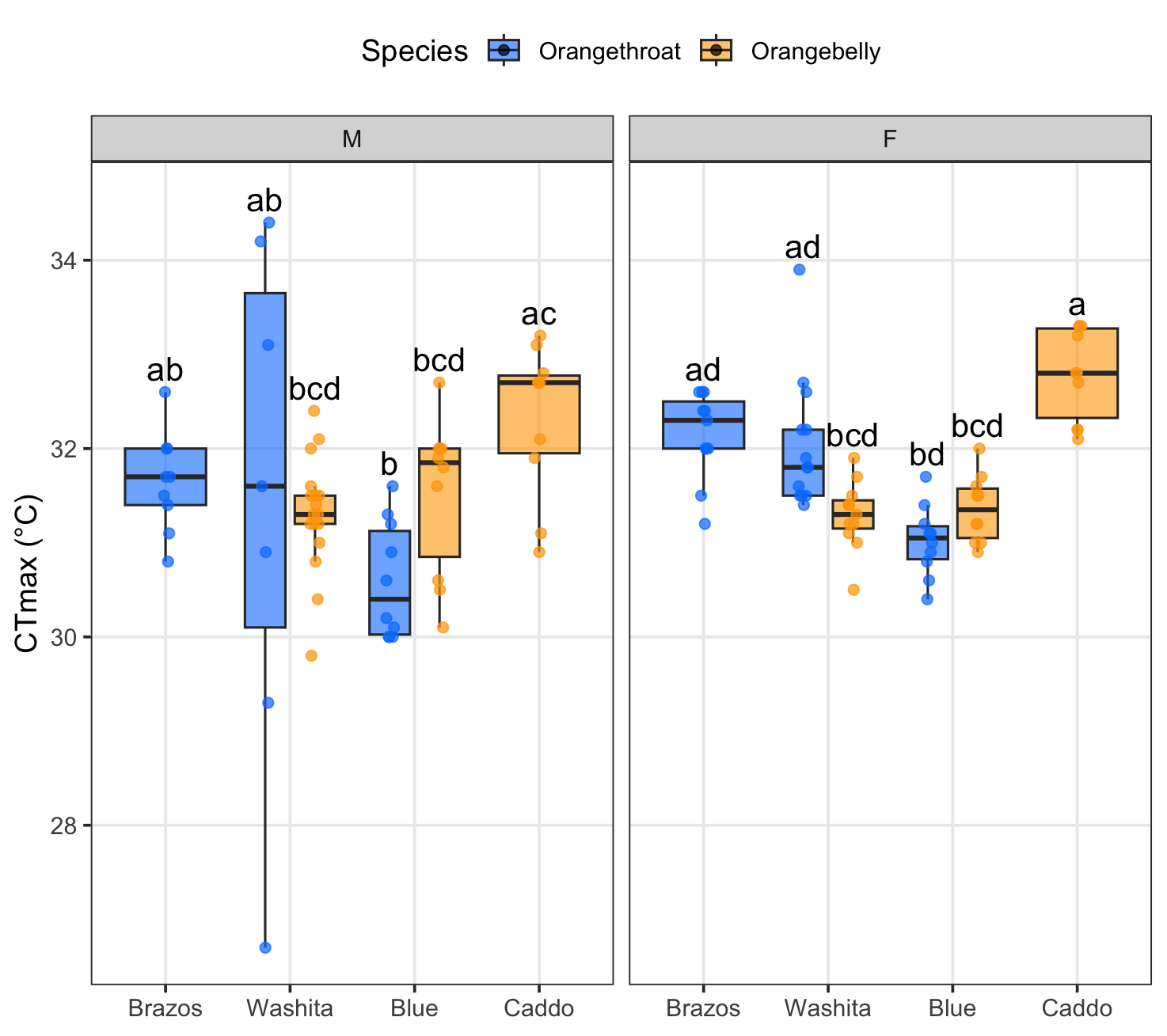

**Figure S1.** CTmax results across drainages and species (*E. radiosum* spp = orange, *E. pulchellum* = blue) split by sex. The single outlier male *E. pulchellum* from Washita is also included here.

**Table S1.** Type III ANOVA for CT_max_ across all four rivers (additive model).

| **Effect** | **df** | **F** | **p-value** |
| --- | --- | --- | --- |
| Species group | 1 | 0.01 | 0.921 |
| River | 3 | 13.65 | <0.0001 *** |
| Sex | 1 | 8.23 | 0.0049 ** |
| Length (centered) | 1 | 5.32 | 0.023 * |

**Table S2.** Logistic regression results for “First Worm Winner” trials (*E. radiosum* spp. success). OR = odds ratio.

| **Predictor** | **OR** | **95% CI** | **χ²** | **df** | **p-value** |
| --- | --- | --- | --- | --- | --- |
| Temp | — | — | 0.32 | 2 | 0.854 |
| River | 0.54 | 0.32–0.93 | 5.04 | 1 | 0.025 * |
| Sex | 0.95 | 0.62–1.45 | 0.07 | 1 | 0.796 |
| Size adv. | 0.66 | 0.39–1.12 | 2.40 | 1 | 0.121 |

**Table S3.** Pairwise species contrasts for overall worm consumption, expressed as odds ratios (OR) for *E. pulchellum*. / *E. radiosum* spp. within river x temperature treatments.

| **Drainage** | **Temp** | **OR** | **95% CI** | **p-value** |
| --- | --- | --- | --- | --- |
| Blue | Cold | 39.96 | ~20–80 | <0.0001 |
| Washita | Cold | 0.74 | 0.36–1.50 | 0.319 |
| Blue | Room | 3.65 | 1.70–7.85 | <0.0001 |
| Washita | Room | 1.80 | 1.02–3.20 | 0.044 * |
| Blue | Hot | 9.20 | 3.36–∞ | <0.0001 |
| Washita | Hot | 0.38 | 0.12–0.72 | 0.0016 ** |
